# The social shape of sperm: Using an integrative machine-learning approach to examine sperm ultrastructure and collective motility

**DOI:** 10.1101/2020.12.05.413120

**Authors:** Kristin A. Hook, Qixin Yang, Leonard Campanello, Wolfgang Losert, Heidi S. Fisher

## Abstract

Sperm are one of the most morphologically diverse cell types in nature, yet they also exhibit remarkable behavioral variation, including the formation of collective groups of cells that swim together for motility or transport through the female reproductive tract. Here we take advantage of the natural variation in sperm traits observed across *Peromyscus* mice to test the hypothesis that the morphology of the rodent sperm head influences their sperm aggregation behavior. Using machine learning and traditional morphometric approaches to quantify and analyze their complex shapes, we show that the elongation of the sperm head is the most distinguishing morphological trait in these rodents and, as predicted, significantly associates with collective sperm movements obtained from *in vitro* observations. We then successfully use neural network analysis to predict the size and proportion of sperm aggregates from sperm head morphology and show that species whose sperm feature relatively wider heads aggregate more often and form larger groups, providing support for the theoretical prediction that an adhesive region around the equatorial region of the sperm head mediates these unique gametic interactions. Together these findings advance our understanding of how even subtle variation in sperm design can drive differences in sperm function and performance.

**Subject Areas:** Evolution, Cellular Biology, Behavior

## 1. Introduction

Despite sharing a common goal to reach and fertilize an ovum, sperm cells have undergone extensive evolutionary modification and are one of the most diverse cell types known [1]. This morphological variation is especially prevalent in mammals, in which sperm cells exhibit tremendous modifications in their compartmentalized and streamlined design [2–4], particularly in the size and shape of the sperm head [2,5,6]. In most eutherian mammals, sperm heads are round or oval, but in rodents they are typically asymmetric and falciform, featuring one or more apical hooks that vary in their length and curvature [3,7–9]. For sperm head designs that are complex, identifying and quantifying informative morphological features and, thus, their functional significance has been an ongoing challenge across taxa [10–14].

There are several hypothesized functional implications for the size and shape of the mammalian sperm head, including sperm hydrodynamics and swimming performance [reviewed in 15–23] as well as collective sperm behavior [24]. Although relatively rare in nature, sperm conjugation occurs when two or more sperm cells unite, often through adhesion at the head, for transport or migration through the female reproductive tract and has evolved multiple times across independent lineages of internally fertilizing species [25–27]. For example, sperm cells form pairs at their asymmetrical sperm heads in the opossum (*Monodelphis domestica*), which is predicted to produce a more streamlined shape and enhance their velocity [28, but see 15]. Moreover, sperm from diving beetles in the family Dytiscidae exhibit striking behavioral polymorphisms among closely related species, which includes the formation of sperm pairs, aggregates, and larger groups (‘rouleaux’) of hundreds to thousands of cells in which their heads stack together like saucers [26]. A cross-species comparison of these beetles revealed that sperm head shape co-evolved with sperm conjugation and showed that species with wider sperm heads more likely to form groups [29], suggesting that sperm head morphology plays a critical role in regulating sperm behavior.

In rodents, sperm cells have been observed to form aggregates within at least five species [8,24,30,31], and their unique falciform head shape has been hypothesized to facilitate these collective behaviors [24]. However, both intra- and interspecies comparisons attempting to link morphological features of the head and group formation have yielded minimal evidential support. Indeed, most rodents produce hooked sperm [3], yet aggregation is relatively rare. For example, a cross-population study in house mice (*Mus musculus*) found that sperm groups were not common and did not form at their apical hook [32]. Similarly, a study in the sandy inland mouse (*Pseudomys hermannsburgensis*), which produces sperm with three hooks, found no evidence of sperm aggregates in the reproductive tracts of mated females [33]. Finally, in a study of 25 muroid species that examined whether the presence of the hook relates to the formation of sperm ‘trains’ found that in all but one species, over 90% of the sperm swam as solitary cells, despite the presence of the apical hook [31]. Together these studies serve to illustrate that the impact of the apical hook may be minimal or secondary to other morphological features or the shape of the sperm head in facilitating sperm aggregation [34].

Among closely-related species of mice in the genus *Peromyscus*, sperm cells exhibit striking morphological and behavior variation and thus offer a unique opportunity to investigate the influence of sperm head shape on sperm aggregating behavior. While most *Peromyscus* species produce sperm with an overall similar morphological design [3, see exceptions within], evidence collected from museum specimens suggests significant interspecific variation in sperm head dimensions across this genus [35]. Moreover, across the genus, all sperm are released as single cells, but some species produce sperm that loosely adhere to one another at their head region, forming sperm aggregates that vary in size [36,37] and in the relatedness of cells that associate with one another [30]. Theoretical models predict that the two-dimensional aspect ratio [36] and three-dimensional shape [34] of sperm heads in the deer mouse (*Peromyscus maniculatus*) facilitate their aggregation. Here we test this prediction across *Peromyscus* species by characterizing the complex and potentially subtle variation in sperm head shape from high resolution images, which we analyze using a combination of manual and automatic morphometric methods, and the size and proportion of aggregated sperm cells using live cell optical imaging. Using two machine learning approaches, we demonstrate that the focal *Peromyscus* species are classified into two distinct classes of head shapes that differ in their elongation and that these differences in sperm head ultrastructure significantly influences the formation of sperm aggregates. Specifically, our findings support the prediction that the sperm head aspect ratio and width strongly associate with collective sperm behaviors within this group of rodents. Our study highlights the utility of using machine learning to examine complex sperm forms, which play a critical role in their function and, ultimately, success.

## 2. Materials and Methods

### Study subjects

We obtained captive *Peromyscus maniculatus bairdii, P. polionotus subgriseus, P. leucopus, P. eremicus*, and *P. californicus insignis* from the Peromyscus Genetic Stock Center at the University of South Carolina, and *P. gossypinus gossypinus* from Dr. Hopi Hoekstra at Harvard University. We conducted this study at the University of Maryland with approval from the Institutional Animal Care and Use Committee (protocol #R-Jul-18-38). To control for variation in life experience, we sampled all available captive *Peromyscus* species and avoided wild-caught specimens; this reduced the number of species we could sample but provided important control over the rearing and testing environments of all study subjects. We obtained sperm samples from sexually mature males and accounted for relatedness among the focal males by assigning siblings a unique ‘Family ID’.

### Quantifying sperm morphology

We fixed sperm with 2.5% glutaraldehyde in 0.1 M cacodylate buffer at 4°C overnight, then washed cells three times in cacodylate rinsing buffer and dehydrated with a graded series of ethanol and Hexamethyldisilazane. We then sputter coated cells in gold/palladium, imaged them on a scanning electron microscope (SEM; Hitachi SU-3500) to obtain five sperm heads per male for each species and additional images of the midpiece region and/or entire flagella (Figure 1B; see details in Supplementary Methods).

**Figure 1.**
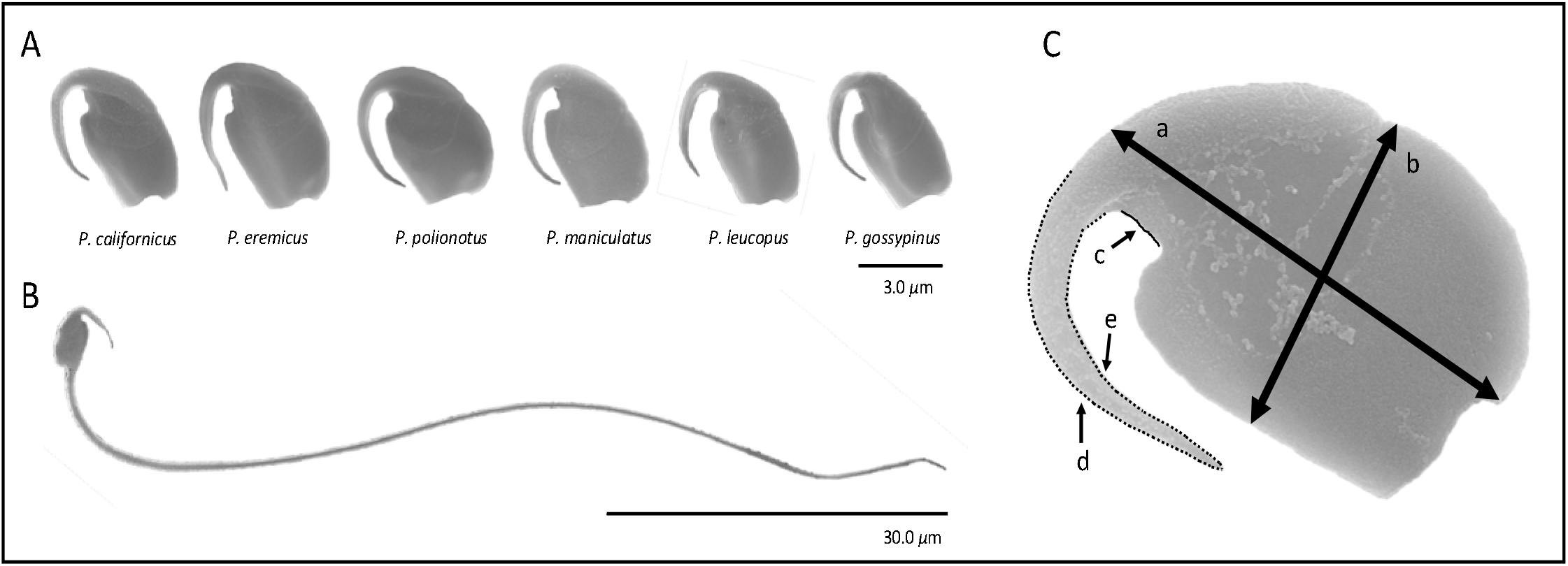
Morphological features of *Peromyscus* sperm. (A) Representative sperm heads for each focal species. (B) An example of a full sperm image. (C) Features measured in manual sperm analysis to estimate sperm head length (a), head width (b), nook length (c), both lower (d) and upper (e) hook lengths.

We manual measured each SEM sperm image using ImageJ (Version 2.0.0, 2017) to estimate sperm features (Figure 1; see details in Supplementary Methods) in triplicate and averaged the values for a single estimate per sperm cell. These data were pooled either across all images per male or across all males per species to characterize morphological traits by males and species, respectively.

### Automatic sperm shape analysis

Anisotropic diffusion [38] was applied to reduce the noise in the SEM images, with integration constant set as 0.1 and κ set as 30, which privileges wide regions over smaller regions. We then detected the boundaries of sperm cells using a rotating derivative of Gaussian filters [39] with the angle interval set as 5 degrees. We applied 72 doG filters with different orientations sized at 31 × 31 pixels on each image, followed by the maximal projection on the stack of filtered images. Before final binarization of max-projected images, we empirically adjusted a threshold to remove noise and closed the holes in images by dilation followed by erosion. To separate the sperm head from the sperm hook for each image, we manually drew a line to cut the binarized image. Finally, we applied the Matlab built-in function ‘regionprops’ to measure the major axis length (i.e., head length), minor axis length (i.e., head width), area, eccentricity (i.e., head aspect ratio) of the sperm head and sperm hook.

### Cross-species comparison via Support Vector Machine

Support vector machine (SVM) finds the best linear boundary to optimize the margin between categories for classification [40]. SVM has been shown to be useful for a wide variety of classification problems, including those in studies of cell morphology [41]. Here we use this method to determine the best linear boundary in terms of classifying the sperm head morphology among species of *Peromyscus* mice with naturally varying sperm aggregation behaviors. We first trained the SVM on the sperm head morphologies of *P. maniculatus* and *P. gossypinus* as measured by traditional manual morphometrics and automatic measurements from high resolution SEM images given that these two species exhibit opposite extremes of sperm aggregation (high and low rates of sperm aggregation, respectively). Measurements included the head length, head width, head aspect ratio, head area, nook length, hook length, hook area, hook width, and hook aspect ratio. Then we selected the most informative features with largest ranked weights in SVM [42], retrained the SVM with the selected features and compared the performances of the retrained machines with the original machines.

To estimate the performance of the SVM, we applied the holdout method [43] in which the dataset was first shuffled then split into training data and test data made up of 70% and 30% of the dataset, respectively. The SVM was trained based on the training dataset, then we calculated the errors of the training dataset and test dataset. We performed 1000 iterations of these holdout processes and then used the average test error to estimate the performance of the SVM. With this estimation method, we validated that the retrained machine with selected features performed similarly to the machine with all features; for both SVMs, training and test errors were low, and the deviation between test errors and training errors were mostly less than 0.02 (Figure S2), indicating that the SVMs were not overfit and were suitable for classification of species. Last, we extended the use of the SVM with the selected features for *P. maniculatus* and *P. gossypinus* to classify the sperm cells of the other focal *Peromyscus* species (*P. californicus, P. eremicus, P. polionotus*, and *P. leucopus*). The machine was calibrated to map the outputs to posterior probabilities by fitting parameters of a sigmoid function [44] prior to classifying all species.

### Quantifying sperm aggregation

We recorded a minimum of five 5-sec videos at 60 frames/sec per male, except in one case we were only able to record four videos using a computer assisted sperm analysis (CASA) system (Ceros II^™^ Animal, Hamilton Thorne). We then manually scored the number of sperm cells present within each track by observing at least three different frames per track, excluding tracks if they involved cells or aggregates that were attached to debris or the slide surface, too out of focus to count the number of cells, interacted with other cells or aggregates (e.g., colliding, joining, separating, or impeding movement), or had a distance average path < 30μm (often repeats of longer, more informative tracks). From the remaining tracks, we calculated (1) ‘aggregate size’ by dividing the number of aggregated sperm cells by the total number of aggregates and (2) the proportion of aggregated cells by dividing the number of sperm cells in aggregate by the total number of sperm cells. We pooled these data across all males per species to characterize sperm aggregate size and proportion by species [45].

### Predicting sperm aggregation from sperm head shape

To predict sperm aggregation behaviors from sperm head morphological features, we applied a fully connected neural network (FCNN). Due to cross-species similarities of sperm head shapes, we ignored classifying by species and combined the data from all six focal species. The network was composed of one input layer, two hidden layers with 5 and 10 nodes, respectively, and one output layer (Figure S3). The mean-square error with regularization was used as the cost function, and Bayesian regularization backpropagation [46,47] was applied to train the neural network. The initial Marquardt adjustment parameter was set as 0.01 and increase and decrease factors were set as 2 and 10, respectively.

To detect the most relevant sperm head morphological features for sperm aggregation, we enumerated all possible combinations of all morphological features, and then fed each combination into the neural network as the input. We based the most informative sperm-head morphological features for predicting aggregation from the combination with the best performance. We applied a seven-fold cross-validation method [48] to estimate the prediction performance by randomly partitioning the dataset into seven complementary subsets, six of which were used as the training dataset and one of which was used as the test dataset. For each partition, the training was repeated 100 times, then the machine with the best performance was determined based on the largest correlation between the observed outputs and the predicted outputs of the test dataset. The final performance estimation of each combination was calculated by averaging the test errors of all the best machines from seven partitions. Finally, the best combination of features with the lowest test error and without overfitting issues was selected to predict aggregation (Figure S3).

### Statistical analyses

For our manual sperm measurements, we performed all statistical analyses and created all visualizations using R version 3.4.2 [49], and prepared all figures using the ‘ggplot2’ package [50]. To compare sperm head morphology across *Peromyscus* species, we used separate linear mixed models for the sperm head aspect ratio and sperm head width with ‘species’ as the explanatory variable and ‘Family ID’ as the random effect; however, we reverted to a linear model (LM) for both responses because the random effect did not significantly contribute to the response variable. We visually inspected diagnostic plots (qqplots and plots of the distribution of the residuals against fitted values) to validate these models. Post-hoc pairwise comparisons were performed using Tukey HSD adjustments for multiple comparisons from the ‘LSmeans’ R package [51].

To examine the relationship between sperm head shape and sperm aggregation, we adopted a phylogenetic generalized least squares (PGLS) approach [52,53]) using the “caper” [54] and “APE” [55] R packages to account for the evolutionary relationships among the *Peromyscus* species in this study. To do so, we used an ultra-metric phylogenetic tree of *Peromyscus* (provided by Dr. Roy Neal Platt II, Texas Biomedical Research Institute), based on sequence variation in the mitochondrial gene, cytochrome B. The species’ relationships within this tree matched those from other previously established phylogenies of *Peromyscus* [56,57]. We used this phylogeny as a covariate in separate regression analyses to investigate the effect of the sperm head aspect ratio, head width, head length, head area, midpiece length, flagella length, and the head-flagella ratio on both aggregate size and frequency. For each of these models, we considered the total number of sperm cells as an explanatory variable and compared models with and without this variable using Akaike information criterion (change in AIC < 2) to determine the best fitting model. Only the best fitting models are reported here.

## 3. Results

### Sperm head morphology

All sperm morphological and behavioral trait values, including the number and percentage of cells aggregated, obtained via manual measurements are reported for each species in Table S1.

We found that sperm head shapes are distinct among the focal species of *Peromyscus*, despite their close evolutionary relationships. Both automatic and manual measurements of sperm head shape indicate that the head width, head aspect ratio, and head area are the most important morphological features for distinguishing between most pairs of species (see Figure S2 for the weights from all combinations of species pairs). In general, the distributions of posterior probabilities were narrow (Figure 2), indicating that the class for each species was well-defined according to the criteria from the trained SVM, which was based only on the data of *P. maniculatus* and *P. gossypinus* to classify the other four *Peromyscus* species. Both manual and automatic measurements revealed similar boundaries based on SVM classification criteria in terms of sperm head morphology: *P. maniculatus, P. eremicus* and *P. polionotus* have similar sperm head morphologies, whereas *P. gossypinus, P. californicus* and *P. leucopus* do not and are themselves more similar. However, *P. polionotus* show larger variance compared to other species, with over 10% of the population classified as being more similar to *P. gossypinus* (Figure 2).

**Figure 2.**
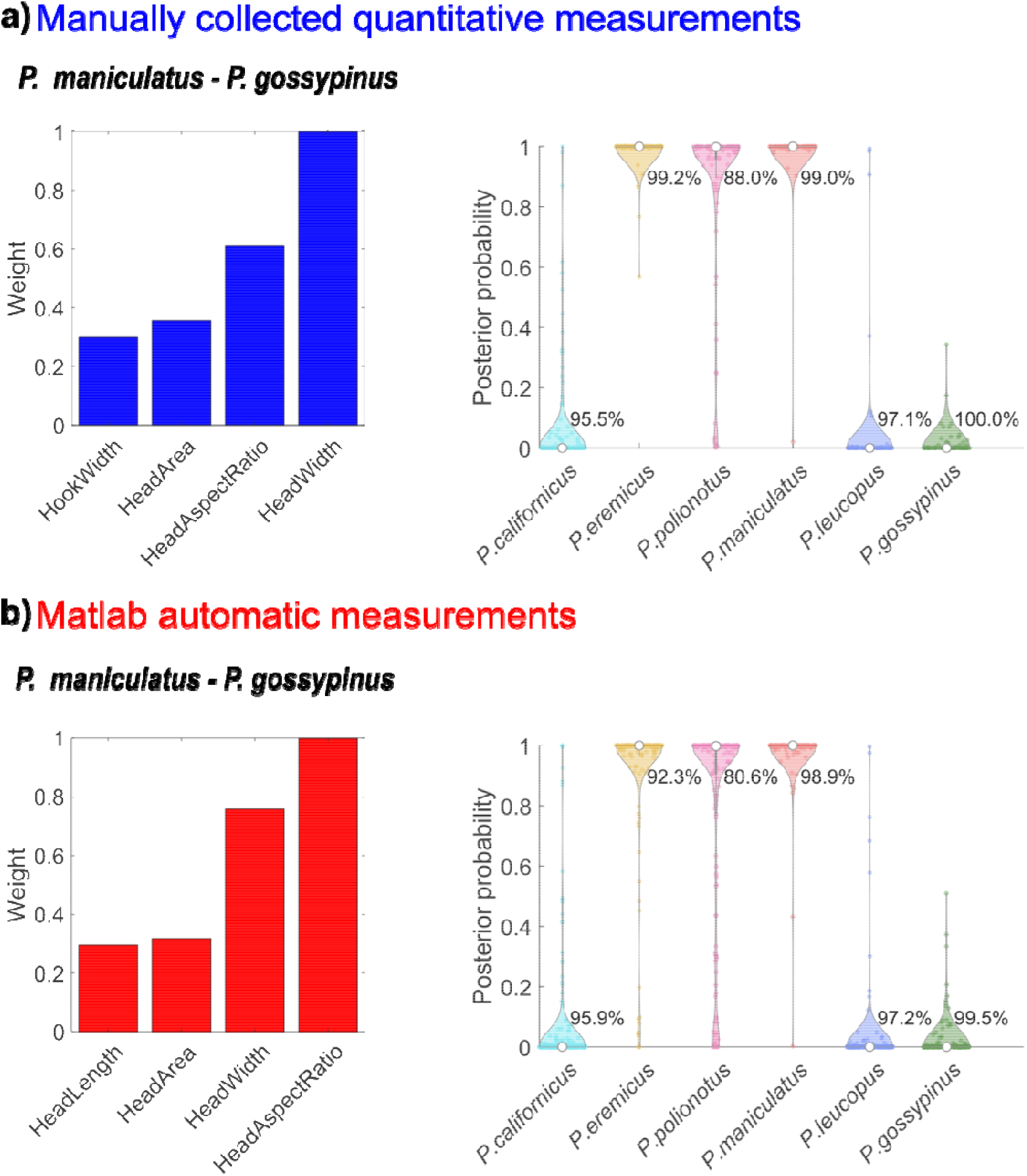
Results from two machine-learning approaches used to identify the most informative sperm head features for distinguishing between six species of *Peromyscus* mice. Analyses using (a) manually collected measurements or (b) measurement resulting from image scanning by Matlab script. The two-class support vector machine (SVM) was trained to classify *P. maniculatus* and *P. gossypinus*, which have high and low rates of sperm aggregation, respectively, based on multiple sperm head features. The SVM was then retrained using only the four morphological measurements with the largest ranked weights (left figures). Based on the retrained SVM for *P. maniculatus* and *P. gossypinus*, posterior probabilities were fitted for all focal species to show how the morphological measurements distinguish among them (right figures). The percentage of samples with posterior probabilities above 0.6 (i.e., more similar to *P. maniculatus*) or below 0.4 (i.e., more similar to *P. gossypinus*) are indicated beside each violin plot. Results from both analyses report that two most informative sperm head features are the head width and head aspect ratio (length/width).

For sperm head aspect ratio, we found more variance across species than within species (s^*2*^ across species = 0.01, s^*2*^ within species < 0.01), and species significantly differed in this trait (LM: F_5,131_ = 91.05, *p* < 0.001; Table 1, Figure 3a). Notably, *P. maniculatus* did not significantly differ from *P. polionotus* (*p* = 0.626). Pairwise comparisons adjusted for multiple comparisons using LSmeans revealed that *P. gossypinus, P. californicus*, and *P. leucopus* all shared similar head aspect ratios (*p* > 0.05 for each comparison), as did *P. polionotus* and *P. eremicus* (*p* = 0.0565).

**TABLE 1.** Fixed effects from a linear model examining the differences in sperm head aspect ratio. All rows are compared with the intercept – *Peromyscus maniculatus*. 95% confidence intervals (CI) were calculated for each effect size.

| Model Term | Beta (SE) | Exp (beta) | 95% CI | t | Pr(>\|t\|) |
| --- | --- | --- | --- | --- | --- |
| Intercept | 1.48 (0.01) |  |  |  |  |
| *P. eremicus* | 0.04 (0.02) | 0.51 | (0.50, 0.52) | 2.16 | **0.0329** |
| *P. gossypinus* | 0.24 (0.02) | 0.56 | (0.55, 0.57) | 12.51 | **< 0.001** |
| *P. californicus* | 0.22 (0.02) | 0.56 | (0.55, 0.56) | 12.83 | **< 0.001** |
| *P. leucopus* | 0.21 (0.02) | 0.55 | (0.54, 0.56) | 10.76 | **< 0.001** |
| *P. polionotus* | -0.01 (0.02) | 0.50 | (0.49, 0.51) | -0.49 | 0.6260 |

**Figure 3.**
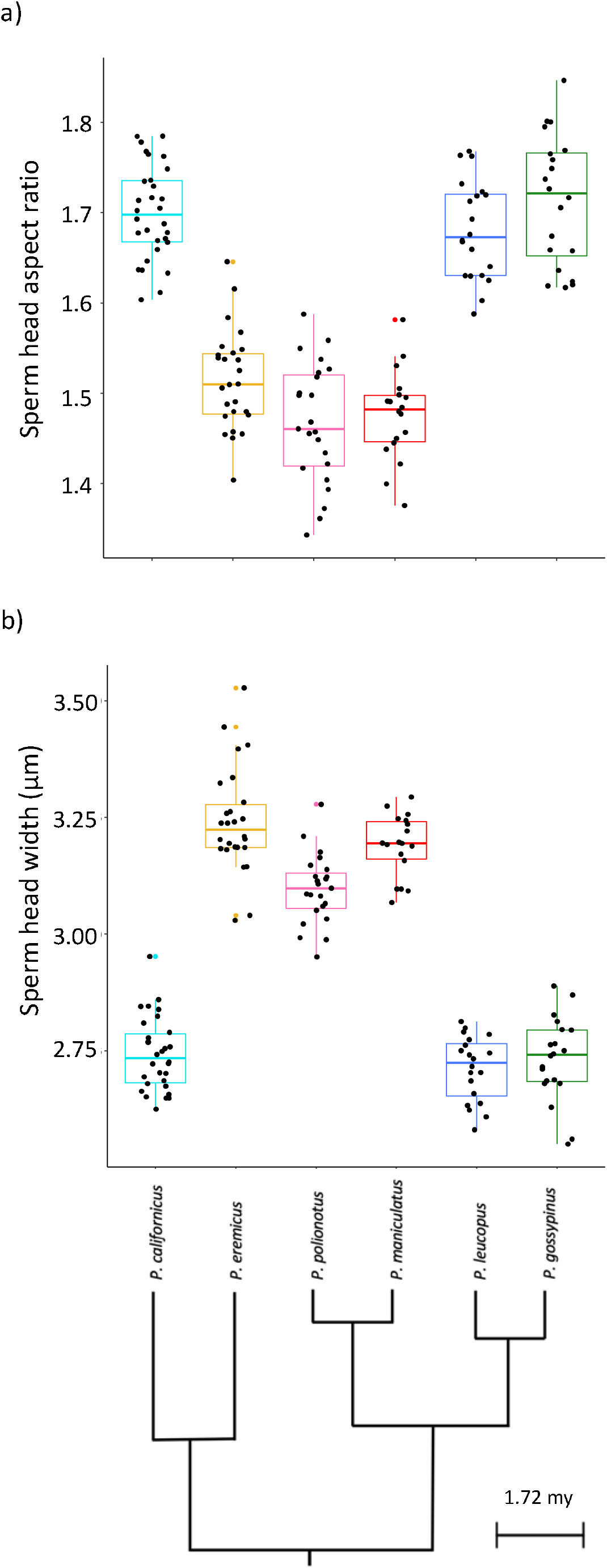
Sperm head morphological variation within and between focal species of *Peromyscus* mice (phylogeny adapted from Bradley et al. 2007) based on manual morphometrics. Box-plots represent median and interquartile ranges of (a) sperm head aspect ratio (length/width), largely driven by differences in (b) sperm head width. Each dot represents mean values per male.

For sperm head width, we found that there was more variance across species than within species (s^*2*^ across species = 0.06, s^*2*^ within species < 0.06) and significant differences among species (LM: F_5,131_ = 199.1, *p* < 0.001; Figure 3b). Pairwise comparisons adjusted for multiple comparisons using LSmeans revealed that *P. eremicus* has the widest cells, *P. maniculatus* has the second widest cells, and *P. polionotus* has the third widest cells, with all other species sharing similarly narrow cells (*p* > 0.05 for each comparison among *P. gossypinus, P. californicus*, and *P. leucopus*).

### Association between sperm aggregation and sperm head shape

Using a fully-connected neural network, we found that among all the combinations of inputs, the network with head width and head area as the sole input provided the best performance to predict sperm aggregation size (Figure S3). These inputs resulted in the smallest test errors and showed no overfitting issues (Figure S3). Moreover, the distributions of errors from the training dataset and test dataset are both centered around zeros and mostly overlapped (Figure 4).

**Figure 4.**
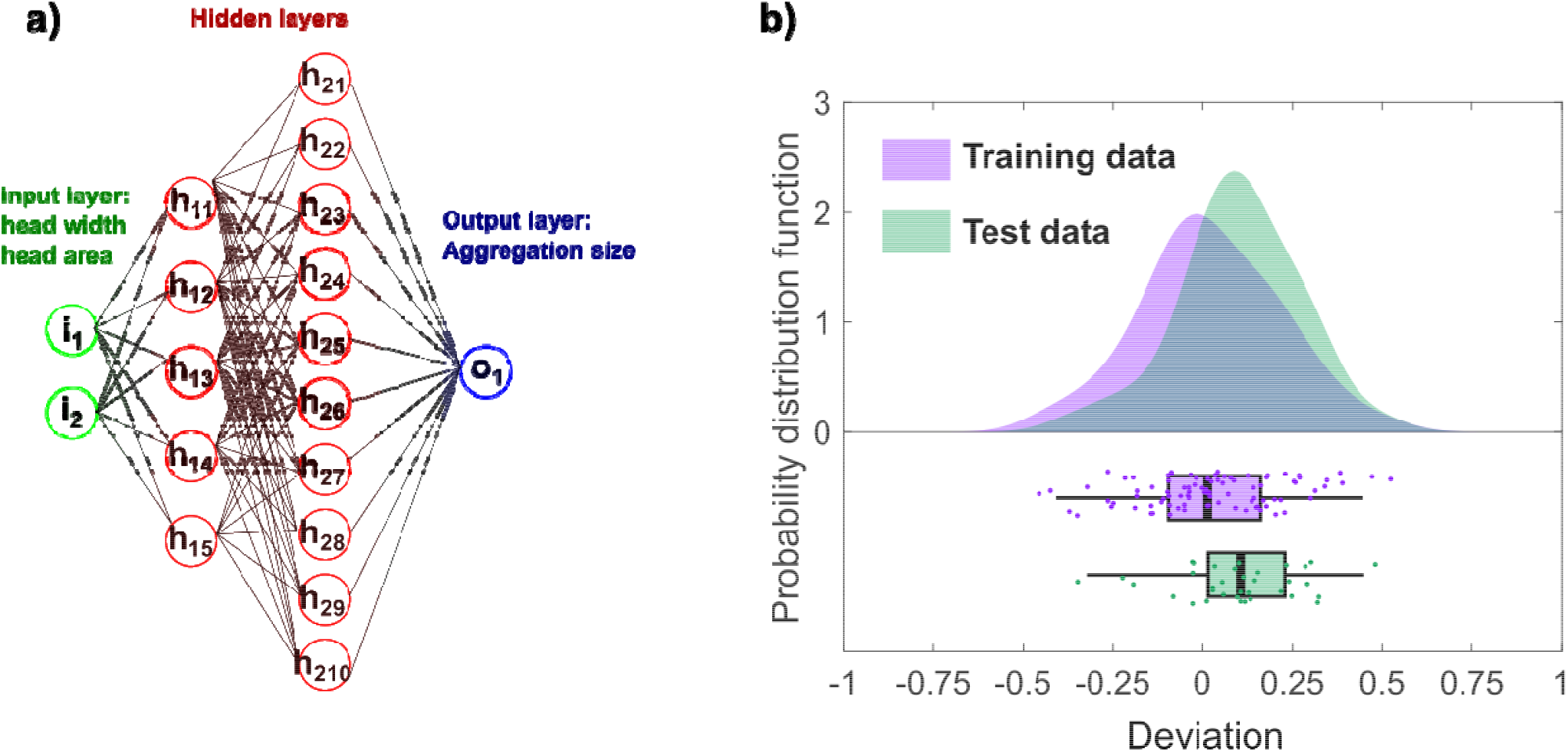
Using a machine learning approach, we predicted the number of sperm cells in aggregate (i.e., sperm aggregate size) from the sperm head morphology of six closely-related species of *Peromyscus* mice. (a) Architecture of the fully connected neural network. (b) Deviation distributions (upper panel) and boxplots (lower panel) of the training dataset (purple) and test dataset (green) generated from our analysis, the latter of which represent median and interquartile ranges. Overlap of the two distributions in the upper panel supports that there is no overfitting, indicating suitable machine performance for predicting sperm aggregation behaviors from sperm head morphology within this group of rodents.

In a complementary approach using statistical models, we found that sperm aggregation, measured by both aggregate size and the proportion of aggregated cells in each sample, were significantly associated with the sperm head aspect ratio. Within our inter-species analysis controlling for phylogenetic relatedness, we found sperm with a smaller head aspect ratio formed larger aggregates (PGLS: F_2,3_= 7.437, *p* = 0.031; Table 2, Figure 5). A similar pattern was observed for head width (PGLS: F_2,3_= 3.617, *p* = 0.076), but such associations were not observed for head length, head area, midpiece length, flagella length, or the head-to-flagella ratio (Table 2, Figure S1). We found similar results for the proportion of cells in aggregate. Within our inter-species analysis controlling for phylogeny, sperm cells within species that had a larger head aspect ratio were significantly more likely to form sperm aggregates (PGLS: F_2,3_= 7.768, *p* = 0.031; Table 2). A similar pattern was observed for head width (PGLS: F_2,3_= 3.269, *p* = 0.086), but such associations were not observed for head length, head area, midpiece length, flagella length, or the head-to-flagella ratio (Table 2, Figure S1).

**TABLE 2.**
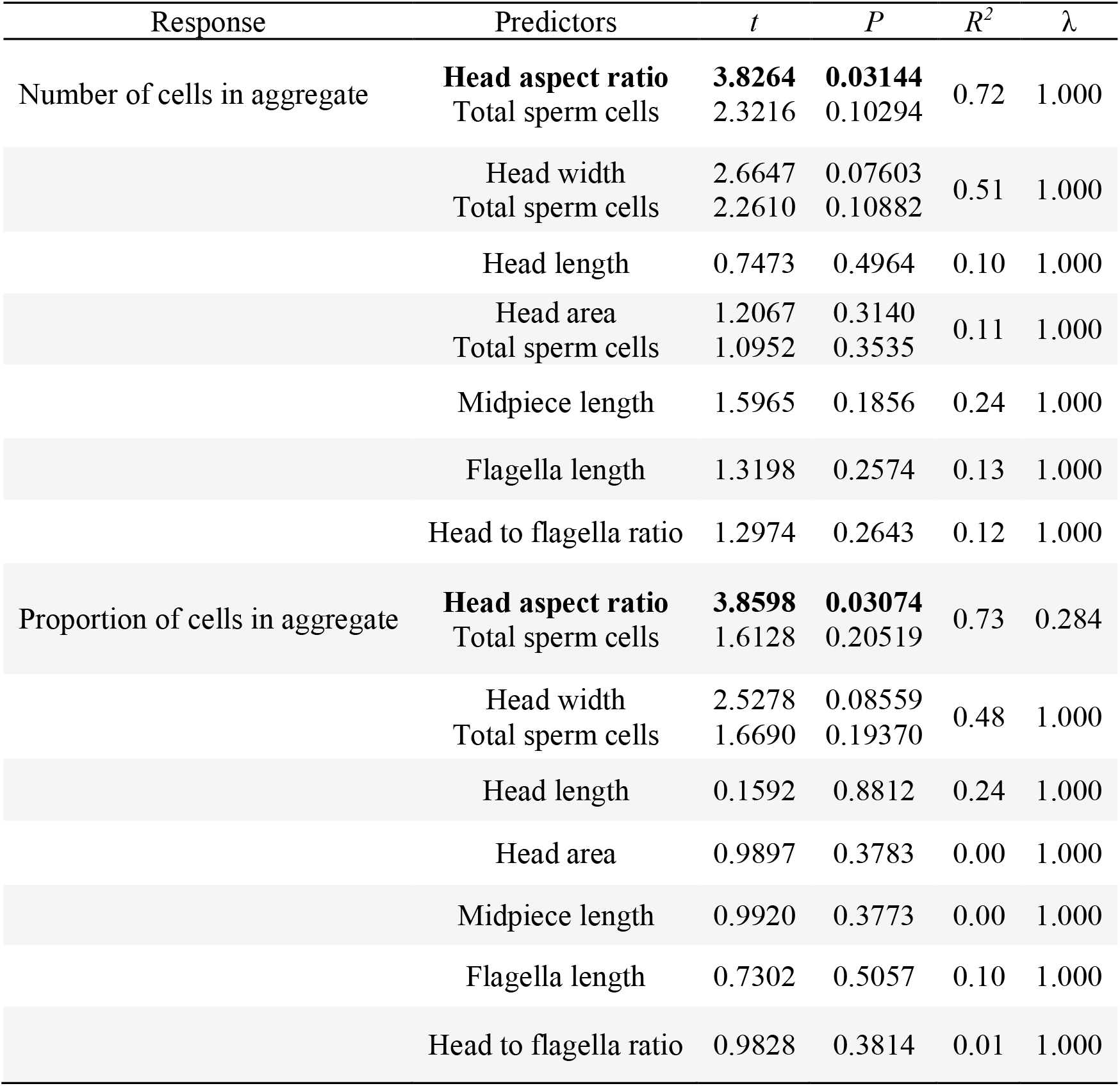
Results from phylogenetic generalized least-squares regression models explaining variation in the number and proportion of aggregated sperm based on morphology. In all models, branch length transformations for lambda, λ, were set using maximum likelihood. Stepwise model simplification comparing models with and without total sperm cells (sperm density) did not change any of these results based on AIC values.

**Figure 5.**
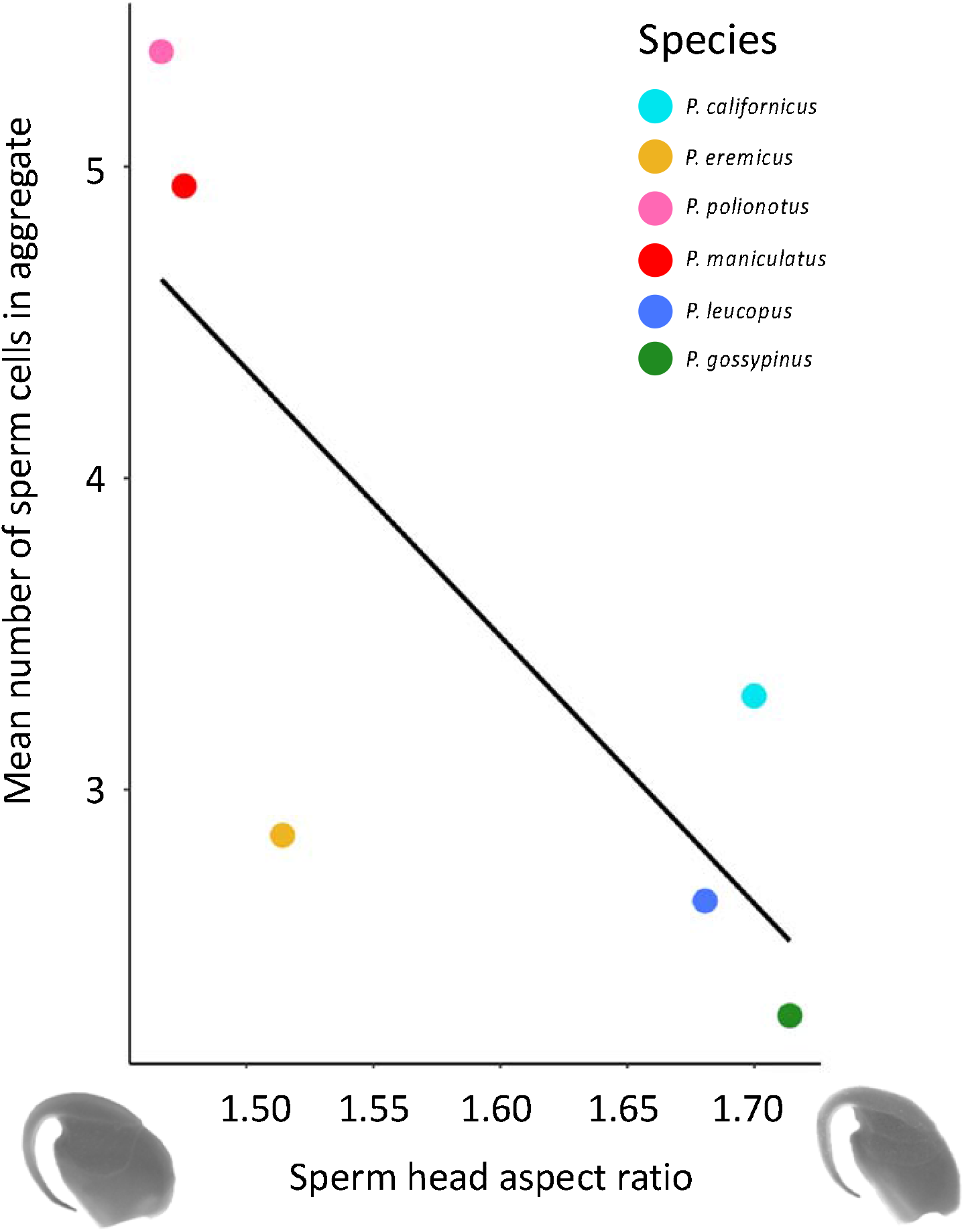
Association between sperm head aspect ratio (length/width) and sperm aggregate size across *Peromyscus* mice, while controlling for species relatedness using a phylogenetic generalized least squares (PGLS). Sperm head aspect ratio negatively correlates with mean number of sperm in an aggregate, such that species that produce sperm with wider heads (i.e., lower head aspect ratio) are more likely to aggregates than species with sperm with elongated heads (i.e., larger head aspect ratio). Dots represents means per species, and species are denoted by distinct colors in the legend above. The line represents the relationship established by the PGLS regression. Note truncated y-axes.

## 4. Discussion

While sperm morphological diversity is well documented and recognized [e.g., 1,12], identifying and quantifying the specific structural attributes that confer functional benefits has proven challenging, particularly for sperm cells with complex shapes [7,11]. Using an integrative methodical and analytical approach that combines traditional morphometry and machine learning, we characterized sperm head shape variation and investigated its role in collective motility in the *Peromyscus* lineage. Our results support the theoretical predictions that sperm with a smaller head aspect ratio (i.e., relatively wider head), are more likely to aggregate and to form larger aggregates when they do [34,36]. Moreover, we find that variation in sperm head dimensions across these captive species is consistent with previously recorded morphological diversity observed from museum samples [35], supporting the use of high-throughput morphometric methods. These analyses reveal that the most distinguishing sperm head trait across our focal *Peromyscus* species is the aspect ratio, and that the width of the cell head, rather than its length, drives the relationship with sperm aggregation behavior. These findings suggest that even subtle modifications in sperm cell architecture can generate large differences in sperm behavior and functionality.

Because rodent sperm are asymmetrical and feature complex head shapes [2,4,8,58], we used a three-pronged approach to characterize and compare morphological variation of sperm produced by *Peromyscus* mice: a traditional morphometric approach using collected-manually measurements of predefined features and two anonymous machine learning methods to computationally identify distinguishing characteristics among our focal species. Importantly, all methods entailed the use of the same high-resolution scanning-electron micrographs of sperm cells produced by multiple males within each species. The results from our machine learning analysis were consistent with those from our statistical analysis of the manually-generated morphometric data, which suggests that machine learning provides a robust and effective method for measuring complex sperm head shapes and can provide similar results to more laborious manual methods. We used a SVM-based analysis of quantitative data collected from sperm head measurements to determine the most informative sperm head features for distinguishing among the focal species used within our study. Our empirical findings indicate that the most useful morphological parameters for classifying the species in our dataset is the aspect ratio and width of the sperm head.

Among the focal *Peromyscus* species within our study, we found more variation in the sperm head aspect ratio and head width across than within species, with some species producing sperm with a relatively wider head (*P. eremicus, P. polionotus*, and *P. maniculatus*) and others producing a more elongated head (*P. californicus, P. leucopus*, and *P. gossypinus*). The elongation of the sperm head, driven by the compactness of the nucleus within [59–64] is expected to reduce hydrodynamic drag and promote more efficient, faster movements [16, but see 15]. Sperm head shape has been observed in both inter- and intra-species analyses across taxonomic groups [17– 19,21–23, but see 65,66]. Given that sperm swimming performance is critical for fertilization success [67,68], variation in sperm head shape that enables greater motility is expected to lead to enhanced fertility potential, as has been shown in humans [69], bulls [70,71], rabbits [72] (Marco et al. 2005), red deer [73], and rams [74,75]), but not boars [76]. Our results suggest another important consequence of sperm head elongation is its influence on the propensity of sperm cells to form collective groups and the size of these formations.

We found that species of *Peromyscus* mice that produce sperm cells with a smaller head aspect ratio, and thus a wider sperm head, form significantly larger and a significantly greater number of sperm aggregates than species that produce sperm with elongated heads. Network analysis of intra- and interspecies data can be used to indicate relationships between morphological features and aggregation. We found that the connection between sperm head aspect ratio indeed was a strong predictor of sperm aggregation in *Peromyscus* mice. Taken together, these analyses provide quantitative support for the hypothesis that collective sperm behaviors are facilitated by the unique sperm head morphological features in rodents [24]. Furthermore, this finding also supports our previous computational model that simulated sperm as self-propelled particles with varying aspect ratios and found that the optimal sperm aggregate size is largely dictated by the geometry of the sperm heads [36].

Our findings in *Peromyscus* are consistent with earlier work on Dytiscid diving beetles indicated a correlated evolution between sperm conjugation and wider-shaped sperm heads [29]. Importantly, if wider sperm heads experience more drag and reduced speeds [16], there may be an evolutionary trade-off between elongated shapes optimized for motility and relatively wider shapes that promote sperm-sperm interactions. Such evolutionary trade-offs between multiple tasks that influence fitness have been observed to drive morphological diversity in bacteria [77] and could potentially explain the benefit of forming groups when cell shape is not optimized for speed. Alternatively, if wider, spherical sperm shapes experience less drag than elongated, prolate shapes when operating in low Reynolds number conditions [15], then selection may favor wider sperm heads overall, but the results from our focal species do not support this prediction.

In *Peromyscus*, the sperm head is the main region of the cell that interacts with another sperm cell when forming with aggregates [36], but we also explored whether sperm flagellar traits are associated with sperm aggregation. Using a subset of males within each species, we found that our focal species varied in midpiece and flagella length as well as the head-to-flagella ratio, consistent with earlier studies of wild-caught mice [35], but we found no significant association between these flagellar traits and sperm aggregation. It has been hypothesized that aggregation enables sperm to collectively increase their length and enhance their motility without the cost of producing an elongated flagellum [29]. A prediction that follows this hypothesis is that species that produce sperm aggregates should have sperm with shorter flagella, but we found no evidence for this prediction in *Peromyscus*, nor was it observed in diving beetles [29]. Furthermore, sperm aggregates that conjoin in a head-to-head orientation, such as in *Peromyscus*, are wider, not longer, than a single cell, which will differentially impact the additive force from the cells and their motility [34,37]. Given the diverse mechanisms by which sperm conjugates form in other taxonomic groups [26,27], including in other rodents in which sperm form ‘trains’ by latching onto the flagella of other cells [8,24], it is likely that there is similar diversity in the morphological attributes of sperm that allow for these unique interactions to occur.

Our finding that the relative width of sperm heads is associated with sperm aggregation may also provide an important clue to localizing and identifying the molecular mechanisms that regulate sperm-sperm adhesion in aggregates. A recent computational study that modeled the sperm head as a three-dimensional, ellipsoid shape equipped with an adhesive band that varied in location and thickness for cell-cell binding revealed that aggregates are more stable and faster when the adhesion is limited to a localized region on the cell [34]. Furthermore, a computer reconstruction of a real sperm head based on morphometric data from the deer mouse (*P. maniculatus*) showed that aggregates with the adhesive region covering only the mid-section of the head, and not the hook, are able to maintain consistently high instantaneous speeds (Pearce et al. 2018). Together, these findings suggest that the geometry of the sperm head and the location of the adhesive region both critically impact aggregate stability and motility and therefore strongly affect the performance of the aggregate [34]. Our results support that the adhesive region is likely to persist as an equatorial band around the sperm head that promotes aggregate formation and stabilization when this region is widened. Moreover, increasing the adhesive range for cellular interactions is predicted to lead to larger aggregates [36], which corroborates our finding that species with wider cell heads also have larger sperm aggregates. Interestingly, the shape of *P. eremicus* sperm heads predicts greater aggregation than we observe; it is possible that despite the width, *P. eremicus* sperm express fewer or different adhesive molecules than their congeners.Although it is beyond the scope of the current study, an important next step is to identify and localize the adhesive molecules that regulate sperm aggregation across these species.

In conclusion, we combine traditional morphometrics with computer-vision algorithms and machine learning to provide insights into the hypothesized relationship between sperm morphology and aggregation by exploiting the natural variation of these features in *Peromyscus* mice. To deal with the challenge of analyzing their complex sperm shapes, we developed an analysis flow to first automatically measure morphologies of sperm cells using traditional computer-vision algorithms and then isolate the most informative morphological features for classifying the sperm heads of the focal species using support vector machine. The automatic analysis proved to be consistent with our manual measurements and successfully predicted sperm aggregation behaviors from sperm head morphologies using the neural network. An association between cell morphologies and function is not unique for sperm cells [41,78–82] and suggests a functional role of the morphology. The methodological approaches we developed in this study further refine that head width and aspect ratio play a functional role more than other features such as the head area. Finally, our integrative approach applies new tools to the study of sperm biology to gain a deeper understanding of how subtle changes in form influence sperm function.

## Acknowledgments

We are grateful to Erica Glasper for providing *P. californicus* males, Hopi Hoekstra for providing *P. gossypinus* males, and the Peromyscus Genetic Stock Center for providing *P. eremicus* males for use in this study; thanks to David Weber and Zaak Walton for help to rear and process the animals used in this study; thanks to Tim Maugel at the Lab for Biological Ultrastructure for training and assistance with SEM and to Bill Breed for assistance with the SEM protocol. We also thank Philip Johnson, Danielle Adams, Sam Church, and Shelby Wilson for statistical advice. We thank Apurva Raghu for helping run the automatic Matlab code to analyze sperm morphologies.

