## Supplementary materials for "The social shape of sperm: Using an integrative machine-learning approach to examine sperm ultrastructure and collective motility"

### Supplementary methods

#### *Sperm collection*

When possible, the focal males used in our experiment were 120 day old virgins, yet we found that *P. eremicus* needed to be >120 days of age and paired with a female to produce sufficient sperm for analysis, and some *P. californicus* males were older and/or paired with a female for a simultaneous experiment (mean ± SE age of total population = 198 ± 12 days). After removing four clear outliers for sperm counts based on our hemocytometer estimates (see methods below), we found an effect of age on sperm counts (LM: $F_{1,117} = 5.92$, $p = 0.017$), but no effect of pairing status (paired $n$ =36, unpaired $n$ =102, unknown $n$ =4) on sperm counts (LM: $F_{1,113} = 1.022$, $p = 0.314$).

To collect live sperm cells, we sacrificed males via cervical dislocation after isoflurane overdose, immediately dissected and made several incisions in the caudal epididymis, and submersed the tissue in 50µl - 1000µl of Modified Sperm Washing Medium (MSWM; 9984, Irvine Scientific) based on epididymal size to standardize cell densities across focal males. We agitated the tissue at 300rpm (ThermoMixer F1.5, Eppendorf) at 37ºC for ten min, inverting the tube at the five- and ten-min mark, and then incubated the tissue undisturbed for two min. Using cut tips with a wider opening, we collected sperm from just below the meniscus to enrich for the most motile cells and continued incubation at 37ºC.

#### *Calculating and adjusting sperm concentration*

At approximately 30 min post-dissection, we assessed sperm cell density within 3ul of each sample, collected from just below the meniscus, in 20µm deep chamber slides (SC- 20-01-04-B, Leja) using a green filter (Hamilton Thorne, product number 720541) to enhance image contrast at 100X magnification with phase contrast (Axio Lab.A1, Zeiss) using a computer assisted sperm analysis (CASA) system (Ceros II™ Animal, Hamilton Thorne). Post-hoc cell density estimates conducted using a Neubauer-improved hemocytometer (220071, Marienfeld Superior) allowed us to confirm that manual sperm counts positively correlate with the CASA estimates (LM: $F_{1,121} = 39.18$, $p < 0.001$). We recorded five 5-sec videos to capture a range of 60-80 cells per video, which we determined was an ideal concentration of cells for CASA tracking efficiency. When necessary, we diluted samples further with warm MSWM until this target concentration was reached to ensure a standard sperm cell density across all focal species (sperm cell density *before* dilutions: LM: $F_{5,118} = 10.19$, $p < 0.001$; sperm cell density *after* dilutions: LM: $F_{5,115} = 1.553$, $p = 0.179$). The resulting final solution of sperm cells for each male then served as the same source of live cells to quantify sperm morphology and aggregation behavior, in which samples for each were collected from the same region of the tube and concurrent in time to control for potential effects driven by differences in either sperm population or the time of collection.

*Quantifying sperm morphology*

To prepare specimens for scanning electron microscope (SEM) imaging, we gently pipetted suspended sperm onto a pre-warmed glass coverslip, incubated them at 37ºC for 15 min, and then fixed cells with 2.5% glutaraldehyde in 0.1 M cacodylate buffer (16537-15, Electron Microscopy Sciences) at 4ºC overnight. Next, we washed cells three times in cacodylate rinsing buffer (21497025, 98% Cacodylic Acid, Acros Organics; 0.1 M with $ddH_20$) for 2 min per wash and then dehydrated with a graded series of ethanol, diluted first with $ddH_2O$ and then Hexamethyldisilazane (HMDS: 98%, 999-97-3, Acros Organics). The $ddH_2O$ dilution series included single washes in 50% ethanol for 5 min, 75% ethanol for 5 min, 90% ethanol for 5 min, 95% ethanol for 5 min, and then three washes in 100% ethanol for 10 min each; the HMDS dilution series included a single wash in 66.6% ethanol for 10 min and then in 50% ethanol for 10 min, followed by three washes in 100% HMDS for 10 min each. We glued coverslips to stubs with carbon adhesive tapes, stored them overnight in a vacuum desiccator, and then sputter coated them in gold/palladium (Balzers Union MED 010 Deposition System) before examining them under a SEM (Hitachi SU-3500). We captured digital images of five sperm heads per male for each species at 15,000 × magnification and an accelerating voltage of 20 kV. We selectively imaged cells with apical hooks that were fully visible (not folded under or over the sperm head) to aid downstream morphological analysis (Figure 1A). Moreover, to determine the importance of the sperm head relative to other sperm morphological features, we collected additional images of the midpiece region and/or entire flagella (Figure 1B) for a subset of focal males using the same procedures above at 7,500 × or 1,600 × magnification, respectively, and an accelerating voltage of 20 kV.

We imported SEM images into ImageJ (Version 2.0.0, 2017) to perform manual morphometrics for each sperm cell. For the sperm head (Figure 1), we used the 'freehand selection' tool to remove artifacts of specimen preparation, including debris and membrane stretching. We measured head width and length by drawing a line using the 'straight line' tool such that the main head body was bisected in both directions, and we then calculated the head aspect ratio by dividing the length by the width. We also calculated the coefficient of variation (CV) for the head aspect ratio and head width within each species using the following formula: (standard deviation/mean) x 100. The CV for the head aspect ratio within each species were as follows: 3.1% *P. californicus,* 3.6% *P. eremicus*, 4.6% *P. polionotus,* 3.4% *P. maniculatus*, 3.3% *P. leucopus,* and 4.2% *P. gossypinus*. The CV for the head width within each species were as follows: 2.8% *P. californicus*, 3.5% *P. eremicus*, 2.4% *P. polionotus*, 2.1% *P. maniculatus*, 2.6% *P. leucopus*, and 3.3% *P. gossypinus*.

We measured head area by adjusting the image threshold in red pixilation (min: 40, max: 100) to three consecutive values such that the cell was fully red in color. We measured the length of the nook (i.e., the straight line between the base of the hook and the beginning of the curvature leading to the horizontal shelf of the sperm head) using the 'freehand line' tool. Then we used the 'freehand selection' tool to draw a straight line up from the nook to divide the hook from the rest of the head. We measured the inner and outer hook lengths using the 'segmented line' tool and then calculated an average of the two values. Using the same method to estimate head area, we then measured hook area. We excluded measurements from our analysis if the cell head was blurry, lysed, or broken or if the tip of the hook was curled or not fully visible. To measure the sperm midpiece and flagella, we used the 'segmented line' tool.

*Quantifying sperm aggregation*

To determine the frequency and size of sperm aggregates, we observed and analyzed live sperm cells using CASA. We collected 4μL of suspended sperm cells from the same samples used for our morphometric analysis and gently reverse pipetted it into 12μL of pre-warmed media on a plastic slide within a 9mm x 0.12mm imaging spacer (Grace Bio-Labs, USA) that we then covered with a plastic cover slip. We chose plastic materials over glass materials to avoid cells sticking to the bottom of the slide, which impacted sperm cell densities (personal observation). We recorded a minimum of five 5-sec videos at 60 frames/sec per male, except in one case we were only able to record four videos. The mean (±SE) duration of time between tissue dissection and video observations was 59 (±1) minutes, which varied slightly depending on how long it took us to properly dilute each sample.

**Supplementary Table**

**TABLE S1.** Summary of morphological and behavioral sperm traits (mean ± SE) across *Peromyscus* mice.

| ***Peromyscus* Species** | **Morphological Traits** | | | | | | | | | | | **Behavioral Traits** | | |
|---|---|---|---|---|---|---|---|---|---|---|---|---|---|---|
| | No. males (No. sperm cells) | Head width (µm) | Head length (µm) | Head area ($\mu m^2$) | Nook length (µm) | Hook length (µm) | Hook area ($\mu m^2$) | No. males (No. sperm cells) | Midpiece length (µm) | No. Males (No. Sperm Cells) | Flagella length (µm) | No. males (No. sperm cells) | No. aggregated cells | % aggregated cells |
| *californicus* | 30 (154) | 2.7 ± 0.01 | 4.7 ± 0.02 | 12.1 ± 0.1 | 0.60 ± 0.01 | 4.22 | 1.46 ± 0.02 | 14 (62) | 15.8 ± 0.1 | 13 (54) | 74.2 ± 0.4 | 29 (8658) | 3.3 ± 0.3 | 32.7 |
| *eremicus* | 26 (131) | 3.2 ± 0.02 | 4.9 ± 0.03 | 15.0 ± 0.1 | 0.66 ± 0.01 | 4.78 | 1.88 ± 0.03 | 15 (72) | 15.9 ± 0.2 | 14 (67) | 77.4 ± 0.6 | 21 (4906) | 2.9 ± 0.1 | 30.2 |
| *polionotus* | 23 (117) | 3.1 ± 0.02 | 4.5 ± 0.04 | 13.1 ± 0.1 | 0.53 ± 0.01 | 4.23 | 1.53 ± 0.02 | 14 (100) | 15.3 ± 0.1 | 13 (56) | 73.6 ± 0.5 | 24 (6360) | 5.3 ± 0.3 | 79.5 |
| *maniculatus* | 18 (96) | 3.2 ± 0.02 | 4.7 ± 0.04 | 13.6 ± 0.1 | 0.53 ± 0.01 | 4.20 | 1.43 ± 0.02 | 13 (65) | 16.2 ± 0.1 | 11 (48) | 78.0 ± 0.5 | 18 (4991) | 4.9 ± 0.2 | 81.3 |
| *leucopus* | 20 (103) | 2.7 ± 0.02 | 4.6 ± 0.04 | 11.7 ± 0.1 | 0.52 ± 0.01 | 4.07 | 1.39 ± 0.02 | 11 (54) | 16.3 ± 0.1 | 11 (49) | 76.6 ± 0.3 | 22 (6341) | 2.6 ± 0.1 | 15.0 |
| *gossypinus* | 20 (105) | 2.7 ± 0.02 | 4.7 ± 0.04 | 11.7 ± 0.1 | 0.54 ± 0.01 | 4.39 | 1.42 ± 0.03 | 11 (61) | 17.6 ± 0.1 | 11 (55) | 83.0 ± 0.4 | 21 (5970) | 2.3 ± 0.0 | 9.6 |

### Supplementary Figures

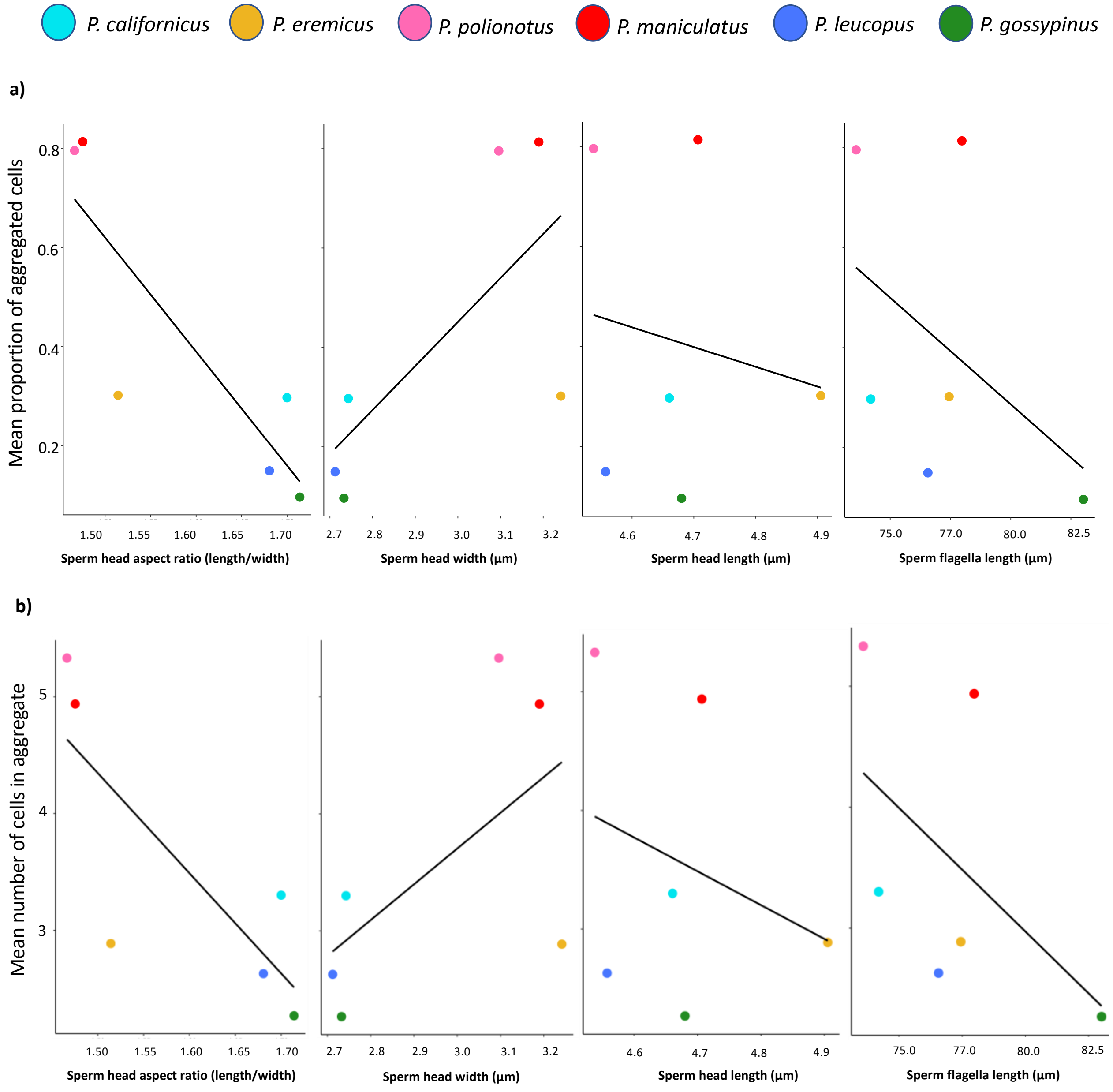


**Figure S1.** The relationships between manually measured sperm morphological features, including the head aspect ratio, head width, head length, and flagella length, and sperm aggregation in six closely-related species of *Peromyscus* mice. Measures of sperm aggregation included the proportion (a) and number (b) of sperm cells that aggregate. When controlling for the close relationships among these focal species using a phylogenetic least squares mean analysis, only the sperm head aspect ratio significantly correlated with the proportion and number of aggregated cells. Dots represents means per species, and species are denoted by distinct colors in the legend above. The black lines represent linear models. Note truncated x- and y-axes.

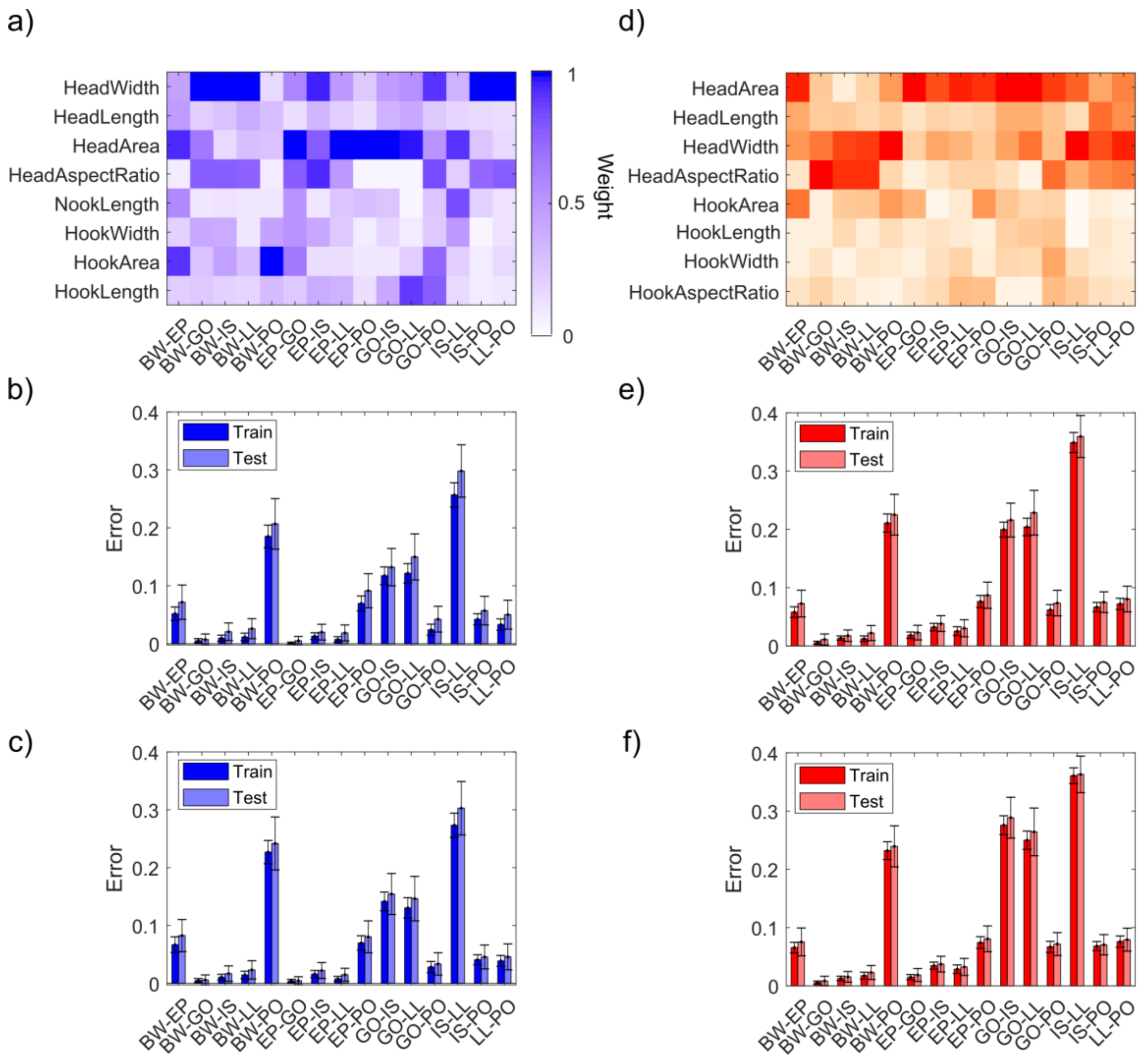


**Figure S2.** Support vector machine (SVM) performances on classifying pairs of *Peromyscus* mice species from manually collected quantitative measurements (panels a-c) and Matlab automatic measurements (panels d-f) based on high resolution SEM images of sperm heads. For each column, the top heatmap shows the ranked weights of various sperm head morphological features for every pairwise species comparison, and the bottom plots show the error rates (with standard deviation error bars) of SVM classification for every pairwise species comparison, either based on all eight head features that were examined (b,e) or the four head features with the greatest relative weights (c,f). The latter analysis yielded similar results to the analysis conducted in panels b and e, indicating that those four head morphological traits adequately distinguished between species of *Peromyscus* mice. Species abbreviations are as follows: *P. californicus* (IS), *P. eremicus* (EP), *P. polionotus* (PO), *P. maniculatus* (BW), *P. leucopus* (LL), and *P. gossypinus* (GO).

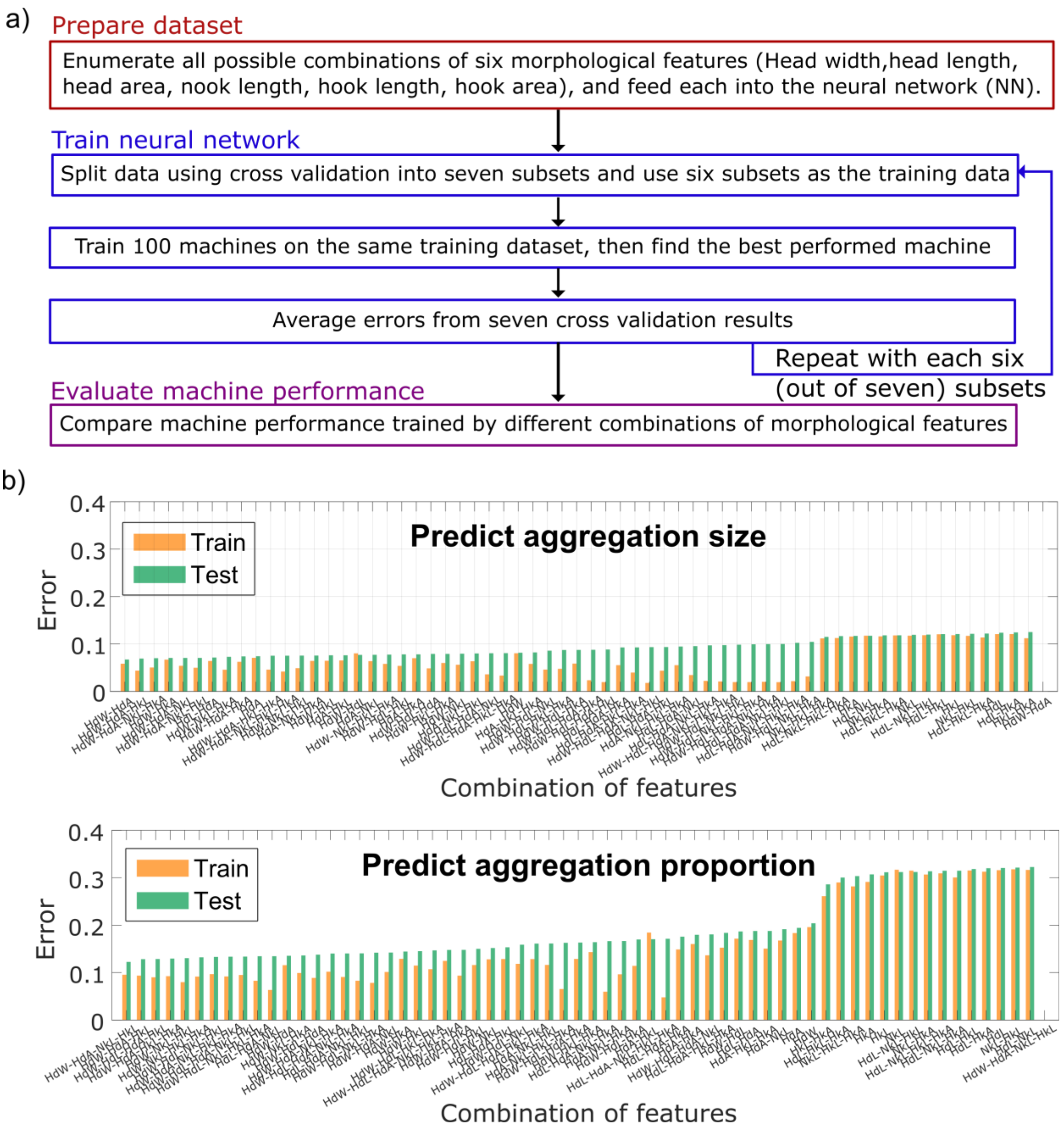


**Figure S3.** We were able to predict sperm cell aggregation behaviors from sperm head morphology across six focal species of *Peromyscus* mice using a fully-connected neural network (FCNN). (a) The analysis flow of selecting the most relevant sperm head morphology features to predict aggregation via FCNN. (b) Performance of FCNN using different combinations of morphology features as the input to the network. Seven-fold cross validation was applied to estimate the network performance, the orange bar corresponds to the average error from the training dataset during CV validation, whereas the green bar shows the average error from the test dataset. The difference between training errors and test errors was small, indicating the trained FCNN did not have severe overfitting issues.
